# Sleep restores an optimal computational regime in cortical networks

**DOI:** 10.1101/2022.12.07.519478

**Authors:** Yifan Xu, Aidan Schneider, Ralf Wessel, Keith B. Hengen

**Affiliations:** Department of Biology, Washington University in Saint Louis; Department of Physics, Washington University in Saint Louis

**Keywords:** Sleep, Criticality, Behavior, Homeostasis, Neurophysiology

## Abstract

**SUMMARY:** Sleep is vitally important for brain function, yet its core restorative process remains an open question. Sleep is assumed to subserve homeostatic processes in the brain because sleep restores functional capacity, and stable function requires compensatory tuning of circuits in the face of experience. However, the set-point around which sleep tunes circuit computation is unknown; for more than four decades, the homeostatic aspect of sleep has been approximated by a heuristic model whose strongest correlate is Slow-wave Activity (SWA). While SWA can indicate sleep pressure, it fails to explain *why* animals need sleep. In contrast, criticality is a computational regime that optimizes information processing capacity, and is a homeostatically regulated set-point in isocortical circuits. Consistent with the effects of waking, criticality is degraded by experience-dependent plasticity. Whether criticality is the computational set-point of sleep is unknown. To address this question, we evaluated the effects of sleep and wake on emergent dynamics in ensembles of cortical neurons recorded continuously for 10-14 d in freely behaving rats. We show that normal waking experience progressively disrupts criticality, and that sleep functions to restore critical dynamics. Criticality is perturbed in a context-dependent manner depending on behavior and environmental variables, and waking experience is causal in driving these effects. The degree of deviation from criticality is robustly predictive of future sleep/wake behavior, more accurate than SWA, behavioral history, and other neural measures. Our results demonstrate that perturbation and recovery of criticality is a network homeostatic mechanism consistent with the core, restorative function of sleep.

## INTRODUCTION

Sleep is believed to restore neural dynamics necessary for complex cognition, sensation, and perception (Cirelli and Tononi, 2008; Ganguly-Fitzgerald et al., 2006; Lim and Dinges, 2010; Lo et al., 2016). This assumes a consistent dynamical structure around which sleep tunes neuronal activity; however this computational regime has not been identified. This problem is complicated by two facts. First, many processes, from extracellular fluid clearance (Kang et al., 2009; Xie et al., 2013) to memory consolidation (Buzsaki, 2015; Diekelmann and Born, 2010) cooccur during sleep, making it challenging to identify which features contribute to the restoration of circuit function. Second, the brain changes at molecular (de Vivo et al., 2017), cellular (Watson et al., 2016), circuit (Halassa et al., 2014), and systems levels (Rothschild et al., 2017) during sleep and wake, further complicating the search. Sleep is proposed to subserve myriad functions, including but not limited to memory (Diekelmann Born, 2010), immune function (Majde and Krueger, 2005), and neuronal physiology (McDermott et al., 2003). Succinctly, a multitude of systems exhibit functional degradation across long periods of wake. However, while disruption in any neurobiological process eventually leads to impairment, it does not hold that every neurobiological process contributes directly to gain control, maximized information transmission, maximal dynamic range, maximal entropy, and robustness to noise, i.e., an optimal computational regime.

Phenomenologically, the factors determining sleep and wake have been described by a heuristic model called the two-process model of sleep regulation (Borbély, 1982; Borbély et al., 2016; Daan et al., 1984). In this model, sleep and wake are regulated by the interaction of a circadian process (Process C) with a homeostatic term that captures the cumulative burden of waking experience (Process S). The two-process model is effective in describing sleep and wake at a behavioral level (Dijk and Czeisler, 1995), as well as some aspects of neurophysiology. Slow-wave activity (SWA) is the well-established, widely-accepted metric of sleep pressure (Franken et al., 2001), and cortical firing rates may similarly fluctuate around locally specified set-points (Thomas et al., 2020). While such measures provide an estimate of the state of the brain, they do not explain how sleep restores circuit function (Borbély, 2001).

The relationship between sleep and an optimal computational regime can be restated as a negative feedback loop in which the difference between optimal circuit tuning and ongoing brain dynamics constitutes an error signal that is minimized by sleeping. If the central purpose of process S is to restore an optimal computational regime, the correctly identified error signal should predict future sleep/wake behavior of the animal. In other words, brains operating near the set-point should be less likely to need sleep in the near future. The ability to provide insight into future sleep/wake behavior is the strongest test of a candidate set-point. In addition to predictive power, the error signal should contain an imprint of recent sleep/wake behavior. Further, if diminution of the error term is truly a core function of sleep, the term should approach a minimum at the end of sleep. Applied to the status quo, SWA can be a powerful indicator of recent sleep history, such as deprivation (Hubbard et al., 2020), but its relationship to future behavior is unclear (Greene and Frank, 2010).

Viable candidates for set-point and error signal must be distinguishable from effectors that contribute to the restoration of function. An optimal state is likely to require many processes that unfold in a sleep-dependent manner, such as the rate of protein synthesis (Adam, 1980; Ramm and Smith, 1990; Seibt et al., 2012). However, while processes such as protein synthesis may contribute to an optimal computational regime, they are not the set-point as they do not intrinsically reflect brain function (O ‘Connor, 1994). In addition, candidates for set-point and error signal must be separable from mechanisms that simply encode time spent in a state. Fortunately, the content of waking can influence the timing and duration of sleep (Abásolo et al., 2015; Dijk et al., 1990; Huber et al., 2004; Milinski et al., 2021), thus allowing the central feature of Process S to be separated from factors that track elapsed time (Tsao et al., 2018). For example, an hour of quiet waking should result in less error around the set-point than an hour of circuit-engaging behavior.

Work in theoretical physics points to a plausible set-point. Diverse networks capable of processing complex information often share key features. Such networks are balanced to avoid runaway gain, encode and transmit information across a range of spatiotemporal scales, and support a broad dynamic range (Beggs and Timme, 2012; Ma et al., 2019; Shew and Plenz, 2013). These properties arise in networks tuned to a nonequilibrium regime of population dynamics called “criticality”, which occurs at a second-order phase transition between order and chaos (Beggs and Plenz, 2003; Cocchi et al., 2017). Near-critical systems exhibit maximized information processing (O ‘Byrne and Jerbi, 2022; Shew et al., 2011, 2009), as well as performance on complex tasks (Cramer et al., 2020).

Criticality has long been hypothesized as a general principle for optimal computation in many domains (e.g., neuronal and artificial neural networks: Langton, 1990; Massobrio et al., 2015), but how it is maintained in the brain remains unaddressed. Criticality is a homeostatic set-point in the visual cortex: monocular deprivation, which drives widespread experience-dependent plasticity (Heynen et al., 2003), initially disrupts criticality which is then recovered over the course of 24 h (Ma et al., 2019). Similarly, daily waking experience can drive widespread changes in diverse circuitry (Abbott and Nelson, 2000; Turrigiano and Nelson, 2004). This raises the question of how circuits maintain robust computational capacity during normal experience. We propose that externally-driven waking experience progressively disrupts the near critical regime and a core function of sleep is to restore criticality. Prior modeling concluded that sleep might pull the brain *away* from criticality (Pearlmutter and Houghton, 2009), while prescient studies of sleep deprivation demonstrated that recovery sleep restores markers of criticality (Meisel et al., 2017b, 2017a, 2013). However, sleep deprivation disrupts nearly all physiological systems (Cirelli, 2006; Longordo et al., 2009; Rechtschaffen et al., 1983). Whether naturally occurring behavior and experience impact the computational regime of the isocortex is an open question.

To understand the role of sleep in maintaining an optimal computational regime in the brain under normal conditions, we monitored circuit activity for extended periods of time, capitalizing on variations in behavior and environment to test fundamental predictions about network set-points. We continuously recorded the spiking of neuronal ensembles in visual cortex for 10-14 d in juvenile rats. Utilizing naturally arising combinations of behavior, circadian time, and environmental conditions, we show that criticality in the visual cortex is systematically perturbed by externally driven experience, and that restoration of criticality is consistent with the core function of sleep.

## RESULTS

To ask whether sleep restores criticality in primary visual cortex of freely behaving rats, we followed extracellular, regular-spiking single unit (RSU) activity continuously for 10-14 d (Figure 1A, 3,325 RSUs total, mean = 35 +/- 7.9 RSUs/12h clustering block). Well isolated RSUs were selected based on waveform properties and spiking statistics (Figure 1B; Hengen et al., 2013; Niell and Stryker, 2008; Buccino et al., 2020). We identified four behavioral states in the data: NREM sleep, REM sleep, active waking, and quiet waking (Figure S1A). Over 10 d, epochs of each behavioral state were evident in both light and dark, and at all zeitgeber times (ZT) within individual animals (Figure 1C, S1C-F). Consistent with previous reports (Watson et al., 2016), there was a significant trend toward lower firing rates in NREM sleep (Figure S1H). All data reported here reflect RSU spiking.

**Figure 1.**
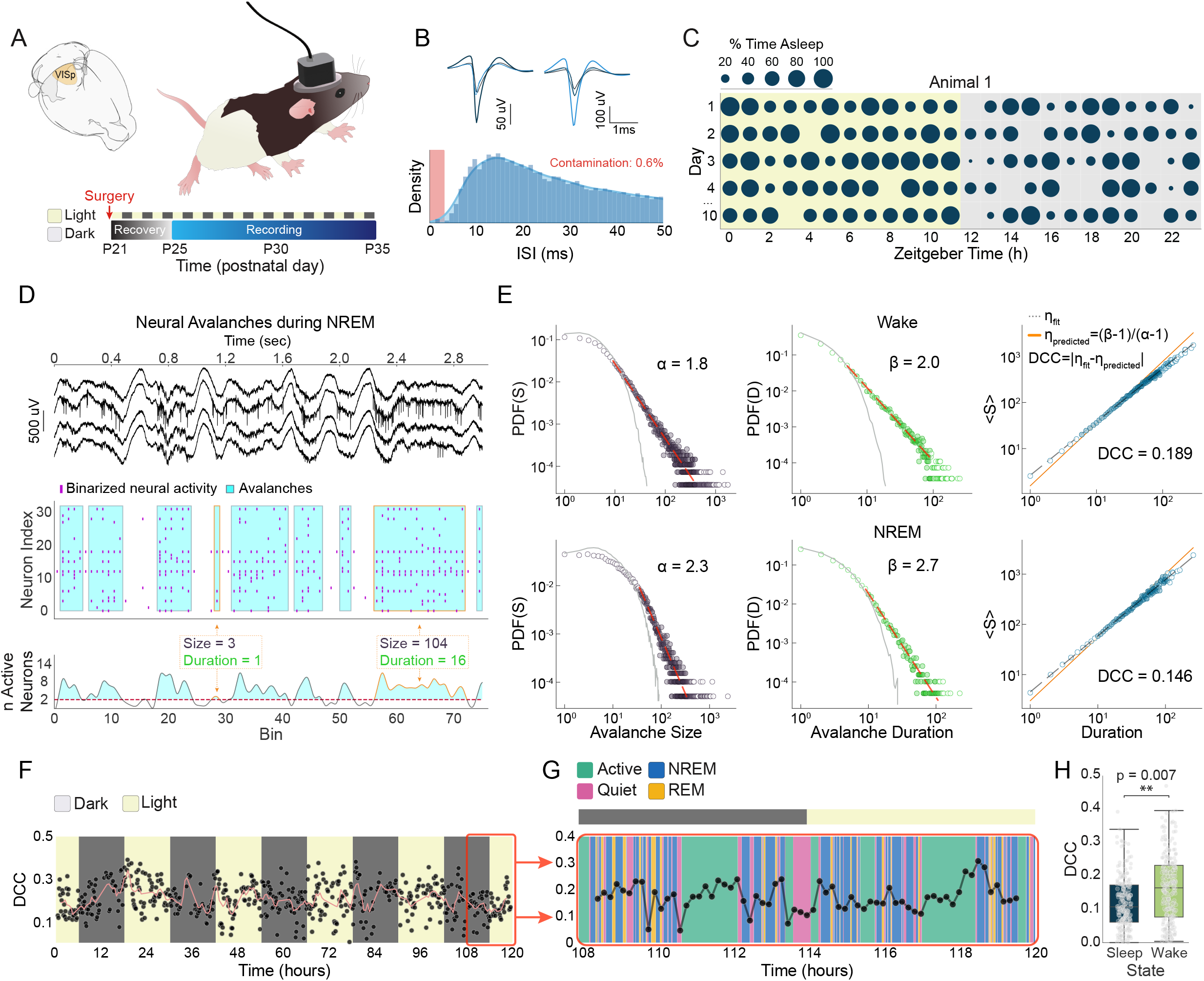
Visual cortical dynamics fluctuate around criticality over long time scales during natural sleep and wake. (A) Experimental approach. Custom arrays were implanted in the primary visual cortex of juvenile rats at P21 and continuous recordings were conducted for >10 d. (B) In addition behavior and local field potentials, single unit spiking was recorded. Two example mean waveforms, each spanning four tetrode channels, are shown (top) as well as an interspike interval histogram (bottom) that reveals a refractory period with low contamination (red box). (C) Individual animals exhibit variability in sleep and wake patterns across the 24 h cycle. Data spanning 5 d from an example animal are shown. The fraction of each hour that is spent asleep is represented by the size of the blue circle. (D,E) Calculation of the deviation from criticality coefficient (DCC) during sleep and wake. (D) Raw data during NREM sleep from four channels (top). Binarized spike counts are extracted (middle), and the integrated network activity (bottom) shows fluctuations. Neuronal “avalanches” start when network activity crosses above a threshold (dashed pink) and stop when it drops below. Avalanches are measured in terms of their size (total number of spiking neurons) and duration. (E) Size and duration distributions could be fitted by power laws in both wake (top) and NREM sleep (bottom). The size and duration distributions scale together and the difference between the empirical scaling relation (blue dots) is compared to the predicted relation. Their difference is DCC. (F) Example animal ‘s DCC over 5 d of light and dark transitions. (G) Zoom on 12 h of data (red box in F) plotted against four behavioral states. (H) Aggregate DCC across 8 animals during wake versus sleep. ** p < 0.01.

We quantified nearness to criticality by first identifying population events, i.e., cascades of spiking activity, often referred to as neuronal avalanches (Beggs and Plenz, 2003). Here, avalanches are periods of time in which population activity rises above a threshold (Ma et al., 2019; Poil et al., 2012; Porta and Copelli, 2019). In critical systems, the probability of observing avalanches of a given size and duration must follow a power law. Further, the two distributions must scale together according to a theory-derived relationship (Friedman et al., 2012; Tang and Bak, 1988; Touboul and Destexhe, 2017). Using the scaling relation, a system ‘s distance from criticality can be quantified by the deviation from criticality coefficient (DCC; Ma et al., 2019). Here, we algorithmically derived the threshold and bin size that maximized the goodness of powerlaw fitting on a per animal basis, separately in sleep and wake (Figure 1D,E, S2A). A 40 ms bin size was uniformly effective across all animals and states. Threshold varied by state. Importantly, thresholds were established in the context of the entire recording and then frozen for all subsequent analyses, such as the evaluation of DCC and its changes over time.

### Near Critical Dynamics Vary by Brain State

Power laws and scaling relations were evident both in NREM and wake (Figure 1D,E, S2A) (REM did not reliably contain enough data to consistently fit power laws). Viewed over days, DCC appeared centered on 0.2 (Figure 1F), consistent with previous evidence of a similar set-point revealed by homeostatic challenge (Ma et al., 2019). However, despite the absence of experimental perturbation, there was noteworthy fluctuation around the set-point. Fluctuations were not a function of circadian time (Figure S2B, p = 0.936, LMER), nor an effect of light or dark (Figure S2C, p = 0.165, LMER). Another possibility is that behavior drove variations in the computational regime of the cortex under normal conditions. On the timescale of hours, variations in DCC appeared to coincide with periods of dense sleep or wake (Figure 1G). Across all animals and the entire 24 h daily cycle, the visual cortical computational regime was, on average, closer to criticality in sleep than in wake (Figure 1H; 0.121 +/- 0.028, 0.162 +/- 0.036, respectively, p = 0.007, LMER).

### Deviation from criticality predicts future behavior

The simplest explanation of the effect of sleep and wake on near-critical dynamics is that DCC covaries with brain state in either a step-wise or time-dependent fashion. In either of these cases, DCC would provide no insight into the neurological impact of waking, which is central to Process S. However, if the maintenance of criticality is the basis of Process S, the error signal (i.e., DCC) should predict future sleep and wake. Crucially, DCC should be dissociable from recent sleep/wake history, as the amount of time spent awake may not indicate the burden of its content (Milinski et al., 2021). To ask whether and to what extent DCC may predict sleep and wake behavior, we first identified unambiguous periods in which animals were predominantly asleep or awake (1h >=66.6% awake or asleep; 193 sleep-dense and 214 wake-dense blocks from 8 animals. Hengen et al., 2016; Vyazovskiy et al., 2009). We then extracted measurements of neural activity in the preceding 2 h window, including DCC, normalized firing rate (FR), and the coefficient of variation (CV) of inter-spike intervals (Figure 2A).

**Figure 2.**
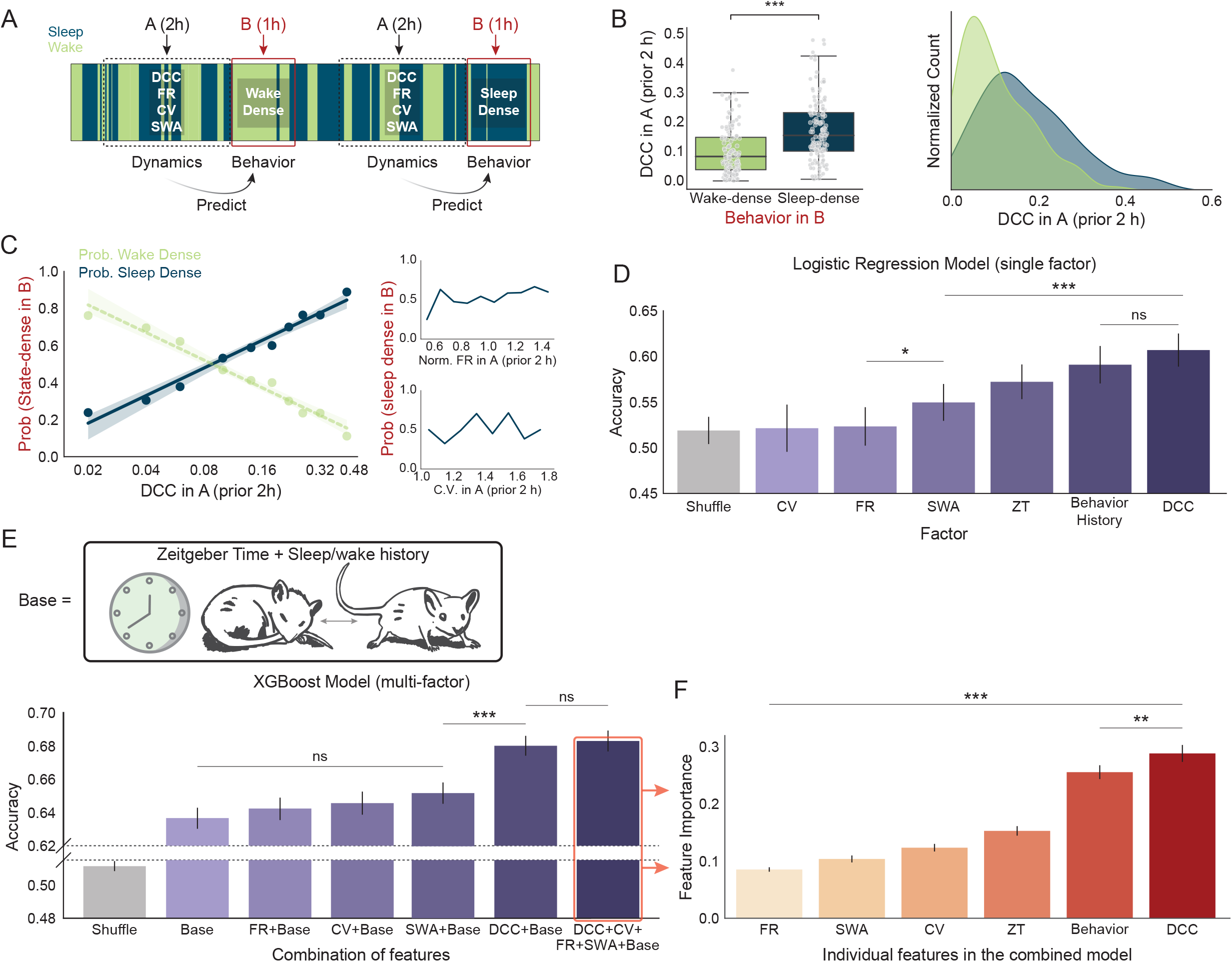
DCC predicts future sleep- and wake-dense epochs better than prior behavior, time of day, SWA, and other neural measures. (A) 1 h sleep- and wake-dense epochs were identified (window B, red) and features of neural activity were extracted from the preceding 2 h window (A, black). (B) DCC in window A is significantly lower before wake-dense than sleep-dense blocks. Box plot (left) and distributions of DCC (right) separated by subsequent behavioral state. (C) (left) The probability of observing a sleep-dense epoch as a function of DCC value. The curve is fitted with logarithmic regression (solid line), and shading indicates 95% confidence interval. Data is from all 8 animals. The probability of correctly predicting a wake-dense epoch is the inverse of the sleep prediction and is shown (green) for ease. DCC is binned such that each interval contains ∼10% of the data. (right) Probability of sleep- and wake-dense epochs as a function of firing rate (FR, top) and coefficient of variation (CV, bottom) measured in the same data as DCC (left). (D) Accuracy of a single-factor logistic regression model in predicting sleep- and wake-dense epochs based on a feature of window A. Gray represents shuffled data/chance. ZT: zeitgeber time, Behavior History: proportion of time spent asleep in window A. (E) Accuracy of a multifactor model (XGBoosted decision tree) trained with combinations of factors. (top) “Base” captures the heuristic Two Process Model: ZT and behavioral history from window A. (bottom) Model performance sorted by increasing accuracy. (F) Attribution analysis of the all-factor model (E, outlined in red). Feature importance is unsupervised and learned by the model and quantifies the weight assigned to each variable to generate the most accurate prediction. ** p < 0.01, *** p<0.001.

Considered across all animals, DCC was significantly lower in the 2 h window before wake-dense epochs than sleep-dense epochs (Figure 2B, 0.102 +/- 0.028 versus 0.173 +/- 0.038, p < 0.0001, LMER). Similarly, the probability of observing a sleep-dense epoch increased monotonically as a function of prior DCC (Figure 2C, p<0.0001, logarithmic regression). The changes in DCC could not be explained by simple changes in neuronal activity: neither FR nor CV from the same data bore a relationship to subsequent behavior (Figure 2C). Together, these data demonstrate that DCC carries generalizable information about the probability of sleep/wake in the near future, consistent with the hypothesis that criticality is a set-point restored by sleep.

Independent of experience, time spent awake influences the probability of future sleep. One possibility is that DCC simply encodes recent sleep history. To test this, we employed a single factor logistic regression model to determine whether individual variables, pooled across all animals, predict subsequent sleep- and wake-dense epochs in a withheld test set (Figure 2D). FR and CV carried no predictive information. SWA, the standard metric of sleep pressure, was 3.1 +/- 2.0% better than chance (p= 0.003). Zeitgeber Time (ZT) and sleep/wake history, which together form the basis of the two-process model, were also significantly above chance (57.2 +/- 1.9%, 59.7 +/- 2.1%; p<0.0001 versus chance), and sleep/wake history carried significantly more predictive information than SWA (p<0.0001). The single variable most predictive of future sleep/wake behavior was DCC (61.1 +/- 1.8% versus chance, p<0.0001. One way ANOVA with a post hoc Tukey test). Across all animals, the predictive power of DCC was significantly greater than all other single factors with the exception of behavioral history (sleep/wake history vs. DCC p = 0.16; all pairwise comparisons p<0.0001).

While these data demonstrate that both DCC and recent behavior carry significant predictive information, they raise the possibility that the information is redundant. While sleep/wake history should inevitably be highly related to the error around the set-point, the error signal should be a more effective predictor if it integrates the neural burden of experience.

To parse this, we trained a series of nonlinear, multifactor models (Chen and Guestrin, 2016) to test how well different combinations of variables predict future sleep and wake. We first approximated the two-process model by combining ZT (Process C) and sleep history (Process S) (Base Model, Figure 2E, top). As expected, the base model performed significantly above chance (Figure 2E, bottom, 63.66 +/- 0.63%, p < 0.0001, One way ANOVA with a post hoc Tukey test). Neither FR nor CV improved the accuracy of the base model (64.23 +/- 0.68%, p = 0.881, 64.56 +/- 0.69%, p = 0.579, respectively, Tukey ‘s test). Surprisingly, the addition of SWA also failed to boost the performance of the base model (65.1 +/- 0.64%, p = 0.15 compared to base). In contrast, DCC significantly improved prediction accuracy (68.01 +/- 0.60%, p < 0.0001 compared to base, Tukey ‘s test). These results suggest that DCC contains unique information about upcoming sleep and wake that is not captured by sleep history and is not reflected in measures of SWA. No further increases in accuracy were evident in a model including FR, CV, and SWA alongside DCC (68.39 +/- 0.63%, p = 0.989 in comparison to base + DCC, Tukey ‘s test).

Given multiple streams of relevant information, data driven models learn to weigh the most reliable variables for a given task. If two variables carry robust information, the model may completely ignore the first if the second subsumes it. To understand the degree to which DCC and behavioral history provide additive, non-overlapping information, we extracted feature importances from the final model containing all variables. FR, SWA, CV, and ZT were of relatively low importance. Behavioral history and DCC were approximately twice as important as ZT, and DCC was slightly but significantly more important than behavioral history (Figure 2F,

28.93 +/- 1.48%, 24.55% +/- 1.21%, respectively, p = 0.0034, One way ANOVA with a post hoc Tukey test). This is consistent with criticality and DCC as the set-point and error signal, respectively, central to Process S and the restorative aspect of sleep.

### DCC in visual cortex increases with time awake in the light and is reduced by sleep

The homeostatic drive to sleep is a product of time spent awake as well as the content of experience (Huber et al., 2004; Kattler et al., 1994; Vyazovskiy et al., 2000). As a result, three predictions about the underlying neural signal should hold: 1) changes in the error signal should be progressive, 2) changes in the error signal should be differentiable from a clock tracking time spent awake/asleep, and 3) the impact of time spent awake should be context dependent. Rodent sleep is highly fragmented, particularly in juveniles (Figure 1C), allowing us to ask whether DCC changes as a function of time, brain state, and circuit input (light/dark) consistent with these predictions.

DCC cannot be calculated on short snippets of data, such as the beginning, middle, and end of a single behavioral epoch. To circumvent this, we divided multi-day recordings into 4 h blocks and concatenated avalanches based on state (sleep and wake). We then regressed DCC within a state against the proportion of the 4 h block spent in that state (Figure 3A). This approach is challenged by the fact that no 4h block comprises pure sleep or wake; each data point thus contains the impact of interleaved states which presumably reduce the signal to noise ratio. However, if the relationship between sleep-need and DCC is strong enough to be detected in this context, there should be a positive correlation between time spent awake and DCC. Further, this relationship should depend on behavior and environment. Consistent with Process S, there was a positive correlation between DCC and time spent awake in the light (Figure 3B; p<0.001, y=0.27x +0.03, LMER). In contrast, waking time in the dark had no correlation with DCC (Figure 3B; p=0.6, y=-0.04x +0.19, LMER), and the distributions of DCC values differed between wake in light vs dark (permutation test, p = 0.007). These data suggest that DCC is a network-level embedding of the cumulative impact of externally-driven activity, a fundamental prediction of Process S.

**Figure 3.**
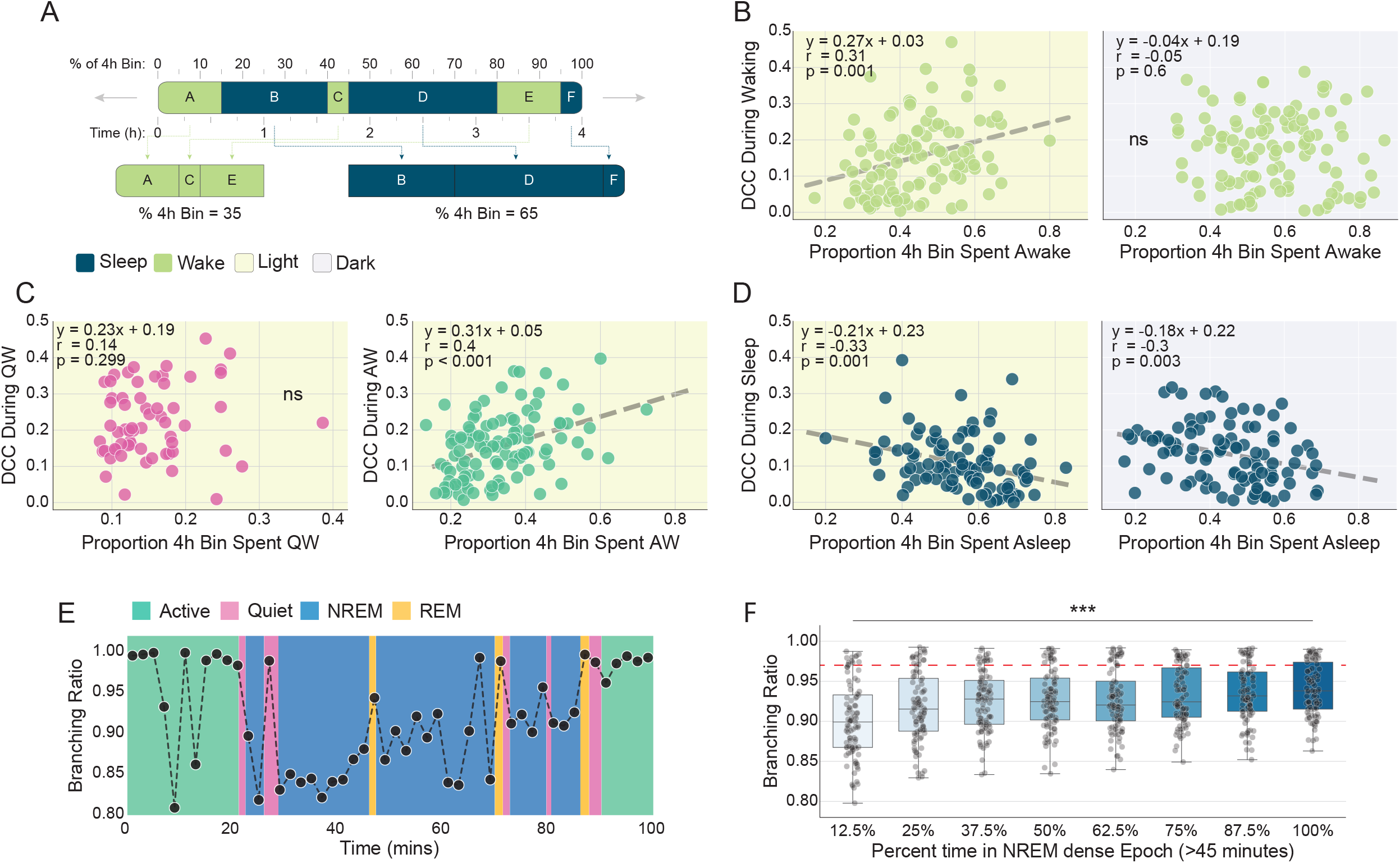
Circuit use during waking undermines criticality which is restored during sleep. (A) To measure progressive changes in DCC as a function of time spent awake or asleep, recordings were divided into 4 h blocks. Each block contained a variable amount of wake and sleep. Avalanches during wake (green) and sleep (navy) were concatenated separately. DCC within a state was regressed against the proportion of the 4 h block occupied by that state. Data in B, C, D, and F are from 8 animals. (B) DCC is positively correlated with time spent in general wake, including quiescence and locomotion, but only in the light (yellow). This effect is absent in dark (gray). Dashed line shows significant linear regression. (C) In the light, time spent in quiescent waking has no relationship with DCC (pink), while the correlation between DCC and time spent engaging in locomotor activity (green) is significant and stronger than general waking in B. (D) In both light and dark, there is a significant negative correlation between time spent in asleep and DCC. (E) Example of branching ratio calculated every 2 min over the course of 100 min. Colors indicate arousal state. (F) Ninety three sleep-dense (>66% NREM) periods of >45 minutes were divided into eighths, and the mean branching ratio was calculated for each division. Red dashed line indicates branching ratio associated with criticality (Ma et al., 2019). *** p < 0.001.

We reasoned that if visual circuit use drives DCC, then the observed correlations should be stronger when constrained to visual behaviors. We divided waking into periods of quiescence and locomotor activity, which is a visual behavior in light conditions (Hoy et al., 2016; Saleem et al., 2013). Compared to all of waking experience, locomotor activity time bore a stronger correlation with DCC (Figure 3C; p<0.001, y=0.31x + 0.05). Consistent with reports that movement engages visual cortical circuitry even in the dark (Guitchounts et al., 2020), locomotor activity in the dark resulted in a weaker but still significant correlation with DCC (Figure S3B; p=0.025, y = 0.14x + 0.08). In contrast, quiescent waking bore no relationship to DCC in either light or dark (Figure 3C and Figure S3B; light: p=0.299, y=0.23x + 0.19; dark: p=0.556, y = -0.16x + 0.25). Together, these data suggest that circuit engagement during wake cumulatively pushes the system away from a near-critical set-point.

Because waking content and time are positively correlated with DCC, sleep must be involved in its diminishment. This could occur in two ways. First, the transition to sleep could involve a stepwise reduction in DCC. Alternatively, sleep could progressively reduce DCC. Consistent with the latter possibility, DCC during sleep significantly negatively correlated with time spent asleep. This is true in both light and dark (Figure 3D; light: p=0.001, y=-0.21x + 0.23; dark: p=0.003, y = -0.18x + 0.22, LMER). To examine the effect of sleep on near-critical dynamics with higher temporal resolution, we calculated the branching ratio, an alternative albeit noisier measurement of criticality that can be estimated in seconds to minutes (Spitzner et al., 2021; Wilting and Priesemann, 2018). We divided sleep-dense epochs into eight bins and calculated the branching ratio within each bin. During NREM sleep, the branching ratio of neural activity progressively increased, converging on criticality at the end of the epoch (Figure 3E, F; p = 0.032, LMER). Alongside the waking data, the recovery of DCC and branching ratio during sleep supports the hypothesis that a core function of sleep is to reassert an optimal computational regime in neuronal networks.

Prior recordings in other isocortical circuits suggest that FR and measures of variance change as a function of time spent awake or asleep (Abásolo et al., 2015; Watson et al., 2016). However, visual cortical neurons may not show such changes (Hengen et al., 2016), allowing for the dissociation of basic spiking properties from emergent dynamics (Ma et al., 2019). In contrast to DCC and branching ratio, CV and FR exhibited no relationship to time spent in either state in any conditions (Figure S3C-F).

Together, these results demonstrate that the value of DCC carries universal information (across animals) about time within a state as well as the content of waking experience. However, these data are correlational in nature.

### Time spent awake is causal in driving the cortex away from criticality

To directly test whether the duration of waking epochs modulates DCC, we used a novel object/gentle handling paradigm (Spano et al., 2019) to extend the length of naturally occurring waking bouts across a 24 h period (Figure 4A; Hengen et al., 2016). To avoid stress caused by cumulative sleep deprivation, we extended every other waking bout to at least 90 minutes, an epoch duration observed approximately 5% of the time under normal conditions. In some cases, animals continued to maintain wakefulness for up to 121 additional minutes (Figure 4B). Animals were then allowed recovery sleep, natural waking, and normal sleep prior to the next 90 min protocol. 90 min of extended waking was sufficient to increase sleep pressure: at the onset of NREM sleep, SWA was increased and then declined over the course of sleep (Figure S4A). In line with the effects of naturally-occuring waking (Figure 3B), we found that experimentally-extended waking significantly increased DCC. DCC returned to baseline levels during the recovery sleep/wake period (Figure 4C,D). Similar results were evident in a branching ratio analysis (Figure 4F; 0.969 +/- 0.005 at start, 0.948 +/- 0.007 at end, p < 0.0001, LMER). However, in contrast to baseline conditions but consistent with our observations of locomotor activity, experimentally-extended waking drove a significant increase in DCC in the dark as well as in the light (Figure 4C; △DCC = 0.105 +/- 0.031, p < 0.0001 light, △DCC = 0.065 +/- 0.023, p = 0.0003 dark). The increase of DCC in dark was marginally less than the increase in light (Figure 4E; p = 0.098, LMER). Consistent with prior research (Hengen et al., 2016), neither FR nor CV changed across the course of extended waking (Figure S4B-E). These results establish that the relationship between waking time and DCC is causal in nature.

**Figure 4.**
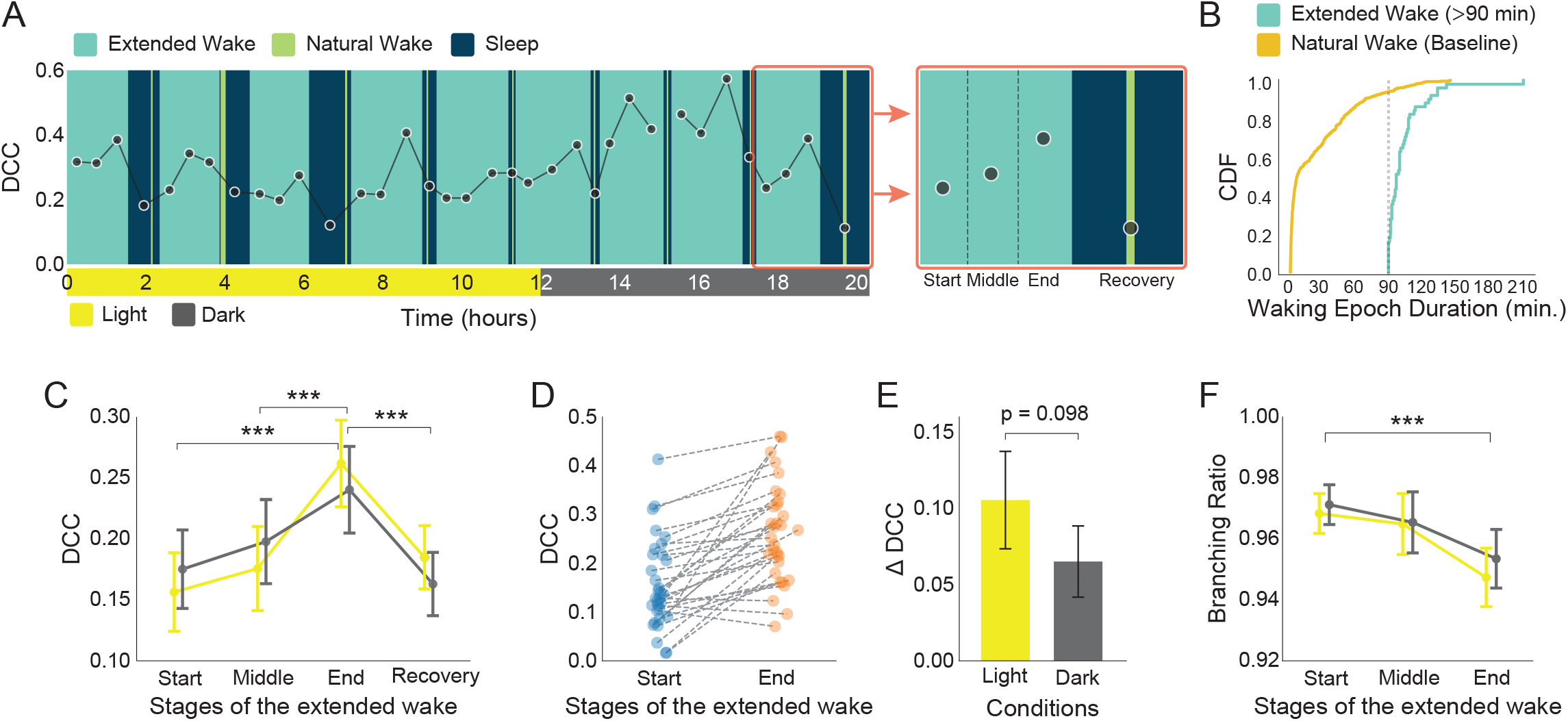
Experimentally-extended waking drives increased DCC consistent with naturally occurring waking. (A) Example of 20 h extended waking protocol and DCC data from one animal. Extended wake bouts (teal, 90 min) are separated by recovery sleep (navy), unperturbed wake (lime), and natural sleep (navy). (Right) each extended waking bout is evenly divided into three epochs (start, middle, and end) in which DCC is calculated. (B) Cumulative histogram of naturally occurring waking epochs under baseline conditions (orange) and extended waking epochs (teal). Experimental intervention ceased at 90 minutes (dashed gray). (C) Summary of DCC changes during all extended waking and recovery periods (n = 8 animals, 55 extended waking epochs) as a function of light and dark (yellow and gray). (D) Individual epoch data (start to end) in light (32 extended waking epochs). (E) The effect of extended waking on DCC is marginally larger in light than dark (p = 0.098). (F) Branching ratio calculated over the same data reveal a progressive departure from 0.97 (the value previously observed in critical models, Ma et al., 2019). *** p < 0.001.

Extended sleep deprivation may drive abnormal processes (Havekes and Aton, 2020; Vecsey et al., 2009). The extended waking paradigm used here did not appear to have pushed animals beyond physiologically normal conditions. First, the effect of extended waking did not accumulate across the 24 h period, consistent with our efforts to avoid a state of sleep deprivation (Figure S4F,G). Second, extended waking had a similar magnitude effect on DCC as 90 min of naturally occurring waking (Figure 3B and Figure 4C). However, increased DCC in the dark suggests that 90 min of experimenter-induced locomotion drove changes in network topology without visual input. In combination with the light/dark effects observed under normal conditions, these results support the conclusion that accumulated activation of visual cortical circuitry is sufficient to disrupt the computational regime of the network.

### Prior behavior predicts nearness to criticality

The theory of Process S proposes that there is hysteresis in its underlying mechanisms. Practically, this means that two equivalent waking epochs can present very different dynamical fingerprints depending on recent sleep history. However, this is challenging to detect, because acute states and behaviors exert complex changes on networks in real time. To test this prediction, we first unified behavior and state as much as possible by algorithmically identifying 193 sleep dense blocks (1h) from all 8 animals. We then regressed DCC during the sleep dense block against the sleep history in the preceding 2 h window (Figure 5A). Despite the fact that DCC was measured over 1 h of sleep in each comparison, there was a significant correlation between DCC during sleep and the recent behavioral history (Figure 5B; p=1*10^−4^, y=-0.17x +0.24, LMER). We observed a nearly identical relationship when examining DCC in wake-dense blocks as a function of recent behavior (Figure S5B; p=2*10^−5^, y=-0.13x +0.21, LMER). In each case, more recent sleep was predictive of lower DCC in the ongoing state. In contrast, prior behavior was predictive of neither FR nor CV during subsequent sleep and wake dense blocks (Figure 5C,D, S5C,D). These data demonstrate that, despite the modulation of DCC by sleep and wake, DCC carries a fingerprint of recent behavioral history. These results suggest that DCC captures the dynamical trace of the recent sleep and wake.

**Figure 5.**
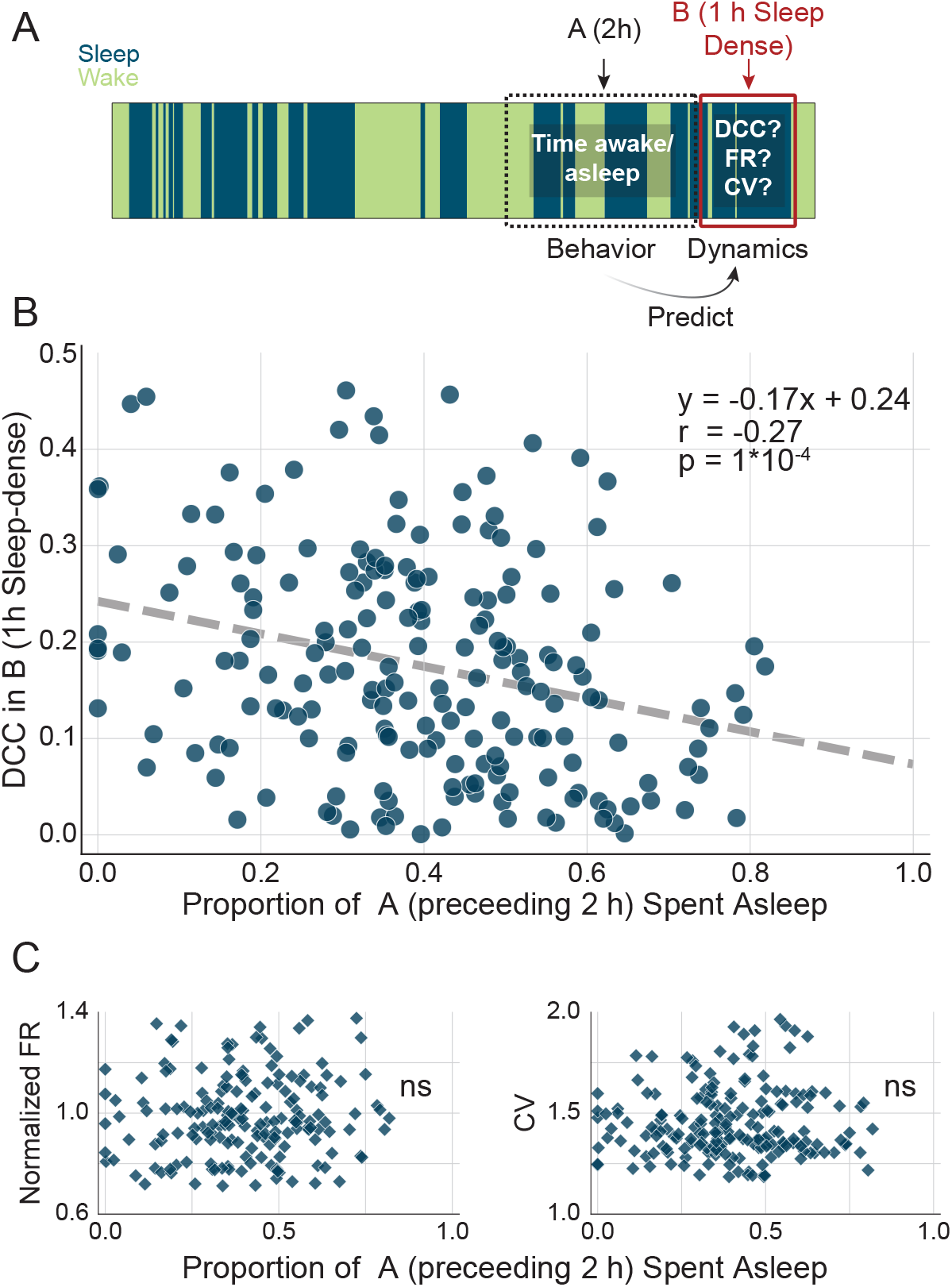
Prior behavior influences DCC within sleep. (A) 1 h sleep-dense epochs are identified (window B, red) in which features of neural activity are calculated. The amount of sleep in the preceding 2 h window (A, black) is calculated as the recent behavioral history. (B) Despite the ongoing effects of sleep on DCC within each sleep-dense block, there is a significant negative correlation between DCC and time spent asleep in the prior 2 h.(C) Analyzed in the same data, there is no relationship between sleep in the 2 h prior and normalized single unit firing rate (left) or coefficient of variation (right) in current sleep dense window.

## DISCUSSION

Previously, we demonstrated that criticality serves as a homeostatic set-point in isocortical circuits in the face of experimental perturbation (Ma et al., 2019). How critical dynamics are regulated in the context of daily experience is unknown. In this study, we evaluated the degree to which the primary visual cortex is near criticality for 10-14 d in freely behaving rats. We examined near-critical dynamics as a function of brain state, behavior, environment, and time of the day in both normal and extended waking conditions. Our data reveal that sleep functions to homeostatically restore the critical regime, which is progressively undermined during waking experience. The extent to which neural dynamics deviate from criticality predicts future sleep/wake behavior and contains information about recent sleep/wake history. This suggests that maintenance of criticality is consistent with the core, regenerative function of sleep. Our results establish a theory-driven, systems-level model describing how sleep and wake modify the computational regime of cortical networks. This model is generalizable across animals and has explanatory power regarding the restorative effect of sleep in the context of daily life.

Neurophysiological hallmarks of criticality are sensitive to brain state and stimulus. External stimuli shift networks towards smaller exponents and longer avalanches (Ponce-Alvarez et al., 2018). On short timescales, strong stimuli can transiently disrupt critical dynamics (Shew et al., 2015), while on longer timescales, the brain broadly operates in a near-critical regime in both sleep and wake (Ribeiro et al., 2010). Sleep and wake states are characterized by slight alterations avalanches statistics, and state transitions drive deviations into subcriticality (Priesemann et al., 2013). These observations suggest that criticality is sensitive to external stimuli and brain states. In other words, this computational regime is highly modifiable and requires active maintenance (Ma et al., 2019). Consistent with this, Meisel et al., demonstrated that sleep deprivation leads to the decay of long-range correlations in both mouse and human brains, an indirect indicator of criticality (Meisel et al., 2017b, 2017a, 2013). These seminal works raised the possibility that sleep can function to restore criticality, but left open the question of direct measurement in the context of normal sleep and wake cycles.

It is likely that a variety of neurophysiological effectors contribute mechanistically to Process S. Slow-wave activity (SWA) is a reliable index of homeostatic sleep pressure (Peter Achermann, 2003) and firing rate deviations around a locally determined set-point reflect Process S (Thomas et al., 2020). Similarly, the frequency of the neuronal “OFF” periods (Vyazovskiy et al., 2009), Lempel-Ziv complexity (Abásolo et al., 2015), and E-I balance (Bridi et al., 2020) are progressively modulated during sleep. However, without prior experimental measurement, it is difficult to predict either the FR (Hz) of an arbitrary neuron or the spectral power (μV^2^/Hz) of a circuit that is optimally functional. Near-critical dynamics are consistent with a theory-derived set-point that predicts behavior across animals and has explanatory power regarding circuit computational capacity. While Process S may be estimated by simply integrating recent sleep/wake history (Borbély, 1982), this does not account for the variable content of waking (Huber et al., 2004; Milinski et al., 2021). Taking a data-driven approach, our present results suggest that DCC conveys the behaviorally-relevant state of isocortical networks more effectively than knowledge of recent sleep history, SWA, or firing rate. Similarly, we observe that DCC in visual cortex is more strongly correlated with time spent in active waking than general waking (including quiescence), and these effects are more pronounced in the light than the dark.

The observation of any relationship between the computational regime of visual cortex and sleep/wake behavior is surprising for three reasons. *First*, Process S presumably integrates the burden imposed by waking across the entire cortex if not brain. Primary visual cortex is a small subset of the brain and is thus likely to contribute only partially to the global signal to sleep. *Second*, Process S-related signals are presumably small under normal conditions, especially in contrast to complete sleep deprivation (Meisel et al., 2017b). This is rationally consistent with experience: animals naturally sleep long before circuit function is severely degraded. In other words, sleep is normally triggered by modest error around the true set point. Apropos this point, SWA is an effective measure of sleep deprivation, but its value in free behavior has been questioned (Hubbard et al., 2020). *Third*, Process S is traditionally queried in the frontal cortices of adult animals. However, we suggest that any network whose local connectivity is affected by experience-dependent Hebbian plasticity will be progressively driven away from a critical topology (Ma et al., 2019). As a result, the primary visual cortex of adolescent animals is an ideal system in which to assess this, as its plastic capacity is enriched (Hensch, 2005) and its input is easily dissociable from behavior (light vs dark). Circuits such as the frontal cortices and hippocampus remain plastic throughout adulthood; we predict that similar patterns will be observable in those networks in older animals.

In combination with our prior work (Ma et al., 2019), our results suggest that the homeostatic regulation of near-critical dynamics plays out in a state-dependent manner under normal conditions. The possibility that the core function of sleep is homeostatic has been long discussed, and recent work highlights synaptic changes that may contribute to the homeostatic function of sleep (Tononi and Cirelli, 2014). Specifically, the Synaptic Homeostasis Hypothesis (SHY) suggests that during waking synaptic weights are globally upregulated by Hebbian processes, which eventually degrade the neuronal signal to noise ratio. Sleep, in contrast, promotes the homeostatic regulation of synaptic strength, e.g., via synaptic scaling, thus restoring the circuit resources necessary for function (Cirelli, 2017). While there is considerable evidence to support such changes (de Vivo et al., 2017; Liu et al., 2010; Vyazovskiy et al., 2009), SHY may be inconsistent with more diverse forms of plasticity during sleep (Benington and Frank, 2003), specifically synaptic strengthening (Aton et al., 2014, 2009). Nonetheless, our results are coherent with the philosophical framework that “sleep is the price the brain pays for plasticity” (Tononi and Cirelli, 2014), although our data suggest that the relevant set-point is implemented at the systems level. This is an important distinction. Rather than tethering waking experience to LTP, we suggest that any combination of experience-dependent synaptic changes are sufficient to erode critical topology. Likewise, no matter the content of waking, sleep must be capable of recovering from the perturbation. As a result, sleep is likely to engage a diverse suite of synaptic and cellular mechanisms necessary to restore an optimal computational regime. Therefore, our study stands to reconcile orthogonal synaptic changes observed during sleep (Frank and Cantera, 2014) by reconsidering the set-point problem at the systems level.

Several limitations of our research merit discussion. Primarily, our recordings are conducted in juvenile rats during the ocular dominance critical period, a developmental time window in which plasticity is enhanced in the visual cortex (Levelt and Hübener, 2012). However, the ability of neurons to exhibit experience-dependent plastic change is present throughout life to varying degrees in different circuits and cell types. If experience-dependent plasticity cumulatively undermines criticality, we predict that the effects described here will be evident throughout life in highly plastic circuits. One effect of this is that specific circuits should play an age-dependent role in driving the need for sleep. A second limitation of this work is that it leaves open the question of how networks restore criticality during sleep. Recent modeling links changes in power law exponents and DCC to the dimensionality and timescales of population dynamics. Specifically, nearness to criticality is influenced by latent variables arising from local activity and external drive, suggesting a locus of control (Morrell et al., 2023). At the cellular/molecular level, plasticity of inhibitory circuits may be foundational to this process (Ma et al., 2019; Stepp et al., 2015; Zeraati et al., 2021), and key molecules that regulate the interaction of interneuron input to pyramidal cells in a sleep/wake-dependent fashion are promising avenues for future work (Pelkey et al., 2015; Severin et al., 2021).

**Supplemental Figure 1.**
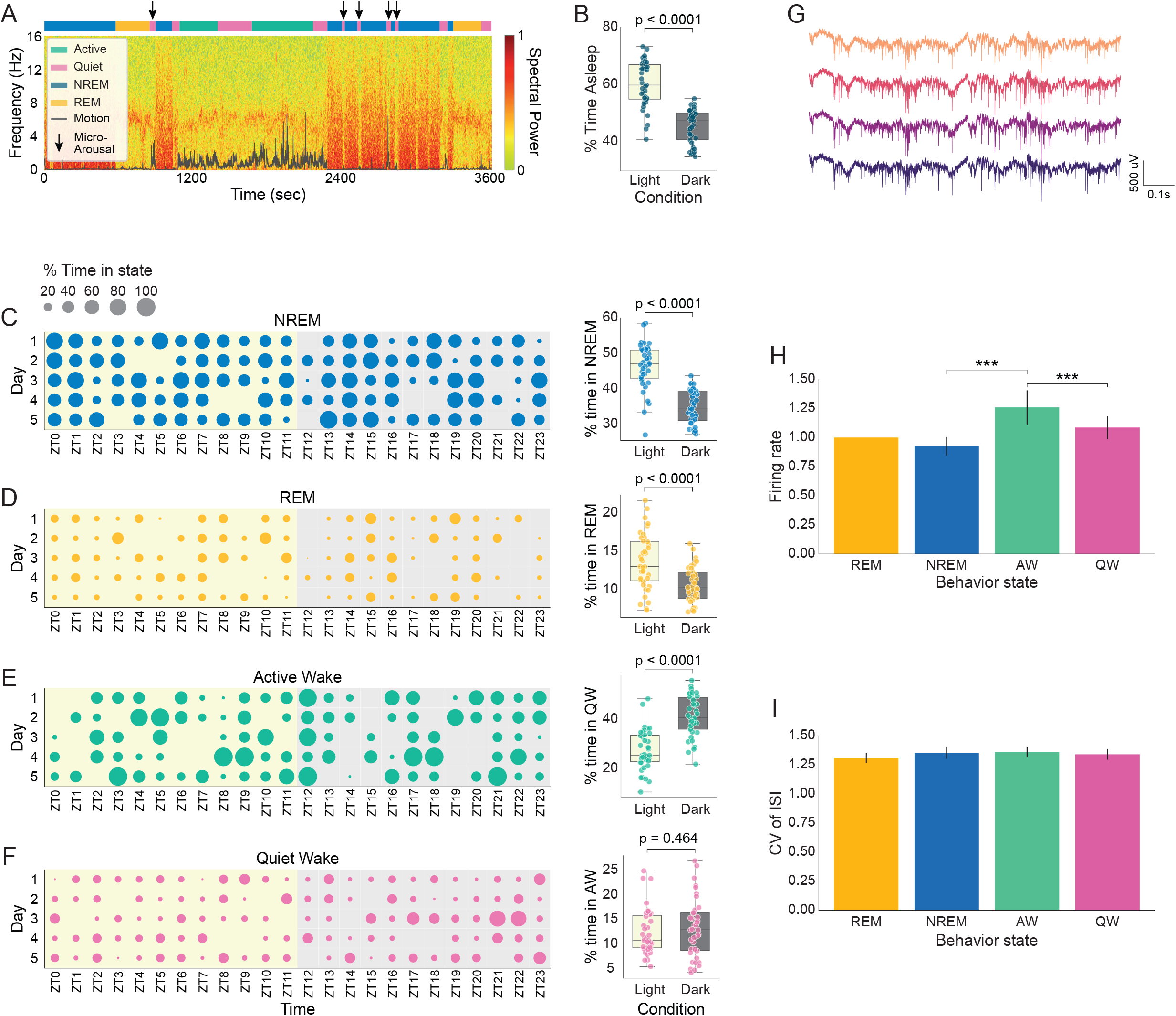
Variability within animals over days in the distribution of wake and sleep substates. (A) Example of sleep scoring shows local field potential recorded across cortical layers (green to red heat map) as well as 15 Hz measurement of motor output (gray). Scoring (semi-supervised) is shown along the top in colored blocks. Black arrows denote microarousals. In this preparation (V1, array of microelectrodes), NREM sleep is characterized by high delta power (0.1-4 Hz), low theta (6-8 Hz), and small muscle movements. REM sleep is characterized by low delta, and obvious theta. Waking is consistently demarcated by low delta and increased motor signal (see Hengen et al., 2016 for details). (B) Young rats spend significantly more time asleep in the light than the dark, although this is highly variable. (C-F) 5 d of data from a single animal showing the percentage of each hour spent in each of four states: NREM (C, blue), REM (D, orange), active wake (E, green), and quiet wake (F, pink). The total time (across 8 animals) in each state as a function of light/dark is shown to the right of each panel. (G) Example raw data traces from a single tetrode (4 channels). Spikes with high signal-to-noise ratio are readily apparent. (H) Normalized single unit firing rate considered by state. Each unit is normalized to its own mean rate during REM. (I) Mean single unit coefficient of variation by state.

**Supplemental Figure 2.**
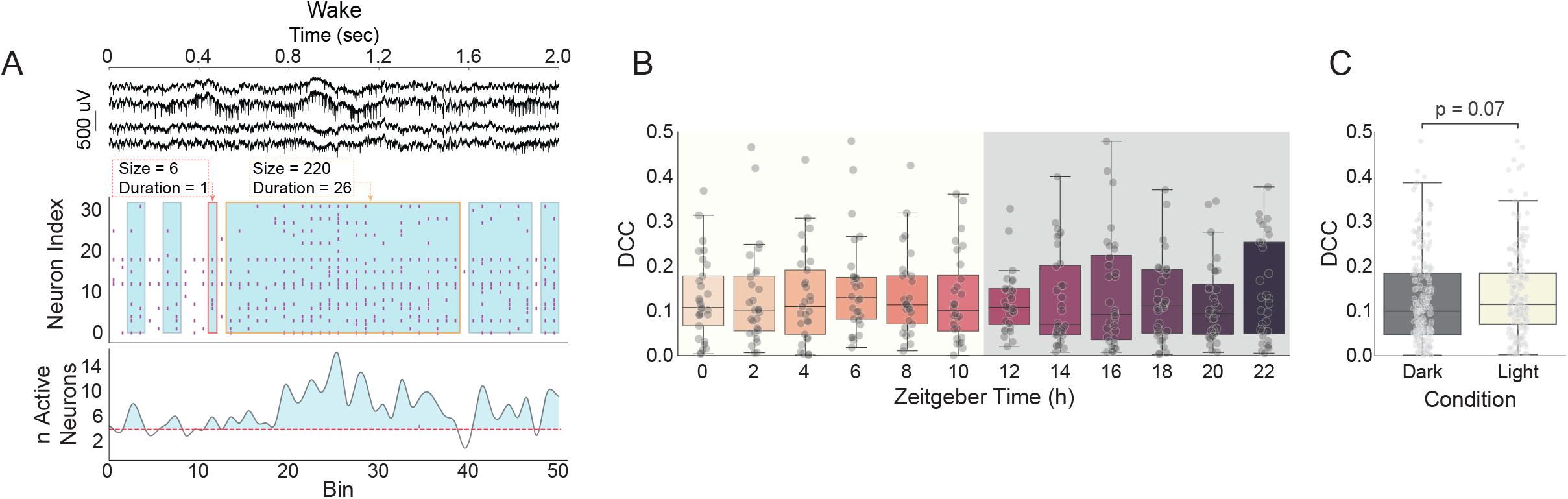
Avalanching during waking and DCC by ZT and light/dark. (A) Raw data during wake sleep from four channels (top). Binarized spike counts are extracted (middle), and the integrated network activity (bottom) shows fluctuations. Neuronal “avalanches” start when network activity crosses above a threshold (dashed pink) and stop when it drops below. Avalanches are measured in terms of their size (total number of spiking neurons) and duration. (B) Variability in DCC is not explained by time of day. DCC data from 8 animals shown in 2 h bins across the 24 h cycle. (C) DCC is not significantly different in light than in dark.

**Supplemental Figure 3.**
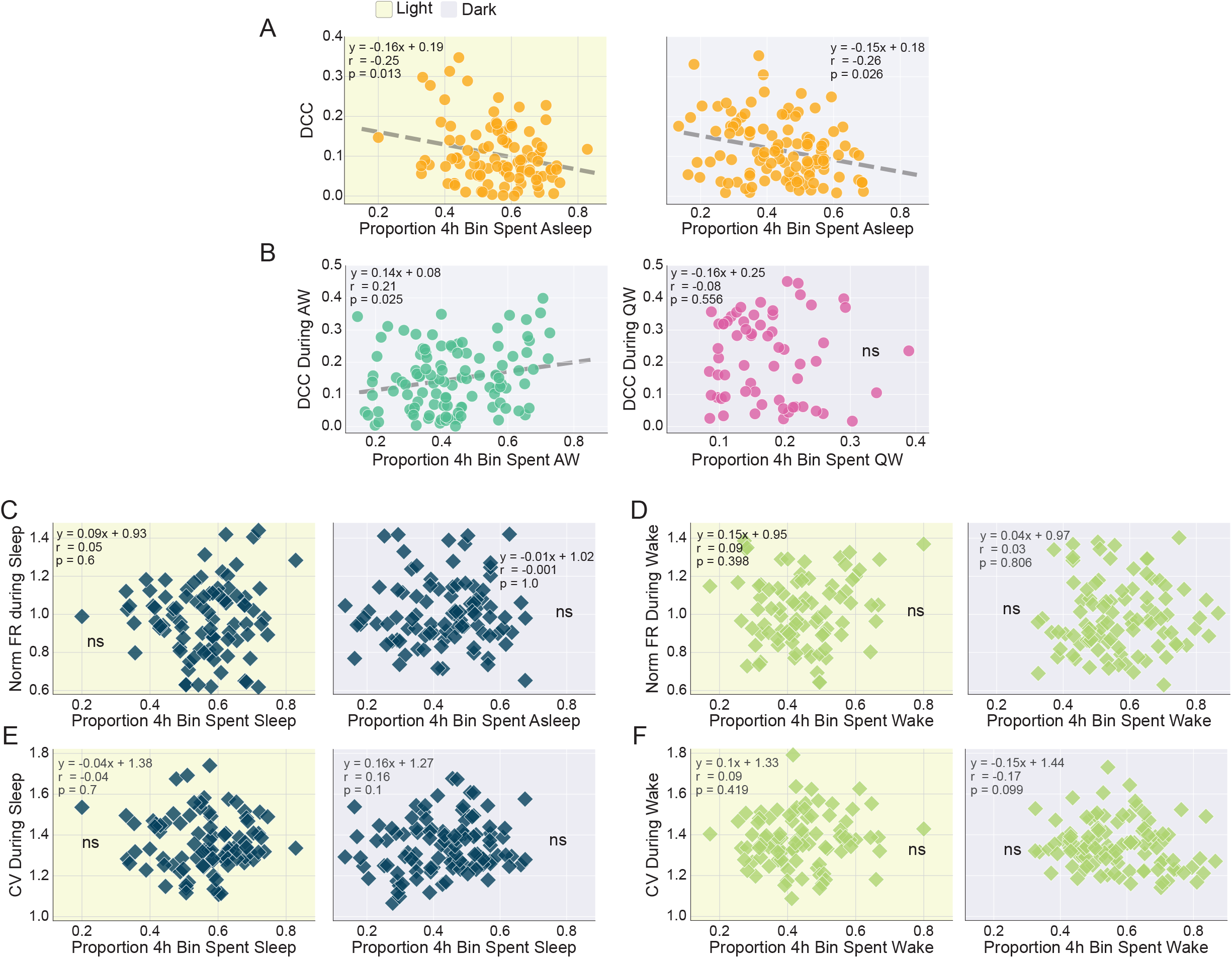
Effects of behavior and environmental conditions on neural dynamics. (A) In both light and dark, there is a significant negative correlation between time spent asleep and DCC (measured across the entire 4h window). Note that this differs from the data in Figure 3D by virtue of examining DCC across the entire 4 h window, inclusive of both sleep and wake. In this approach, each point is 4h of data, but contains mixed states. In Figure 3D, DCC is calculated only during the subset of the window that is spent asleep, and thus has variable time but constant state. The remaining measurements (B-F) are calculated only within a given state, consistent with Figure 3. (B) Time spent in active wake (locomotion) in the dark significantly correlates with DCC. Time spent in quiet waking in the dark has no correlation with DCC. (C) Time spent asleep has no correlation with normalized firing rates during sleep in either light or dark. (D) Similarly, time spent awake has no correlation with normalized firing rates during waking in light or dark. (E,F) Same as C and D but coefficient of variation (CV) of interspike intervals.

**Supplemental Figure 4.**
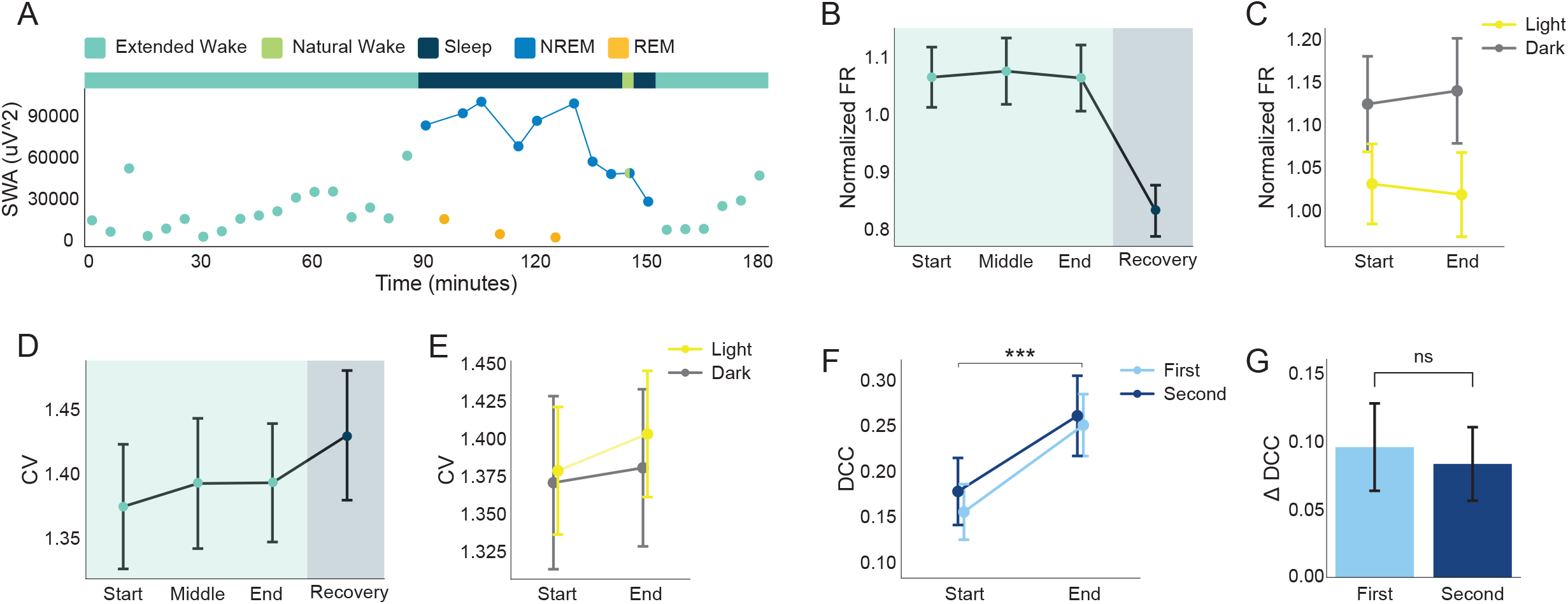
Extended waking data. (A) Process S is traditionally quantified by changes in slow-wave activity (SWA), especially in the context of sleep deprivation (Franken et al., 2001). To confirm that our extended waking protocol is consistent with prior work, we quantified SWA (absolute power) throughout an extended waking epoch and the recovery period. 90 min of extended wake (teal) is sufficient to drive elevated SWA at the onset of NREM (blue) during recovery sleep. SWA progressively declines during NREM sleep. SWA was calculated in 5 mins bins (with a median filter of 4 s sliding windows in each epoch, similar to Franken et al., 2001). (B) Normalized firing rate (FR) does not vary as a function of time spent in extended waking. FR is significantly lower in the recovery phase, the majority of which is NREM sleep (consistent with Figure S1H). (C) FR at the start and end of extended waking, divided by light and dark phases. (D) The interspike interval coefficient of variation (CV) does not vary significantly across the extended waking protocol or recovery period. (E) Same as in C but for CV. (F) The impact of extended waking on DCC is a significant increase between the start and end of the protocol. The magnitude of this change does not differ when comparing the first half of the 24 h period (light blue) to the second half (dark blue). (G) Quantification of data in F.

**Supplemental Figure 5.**
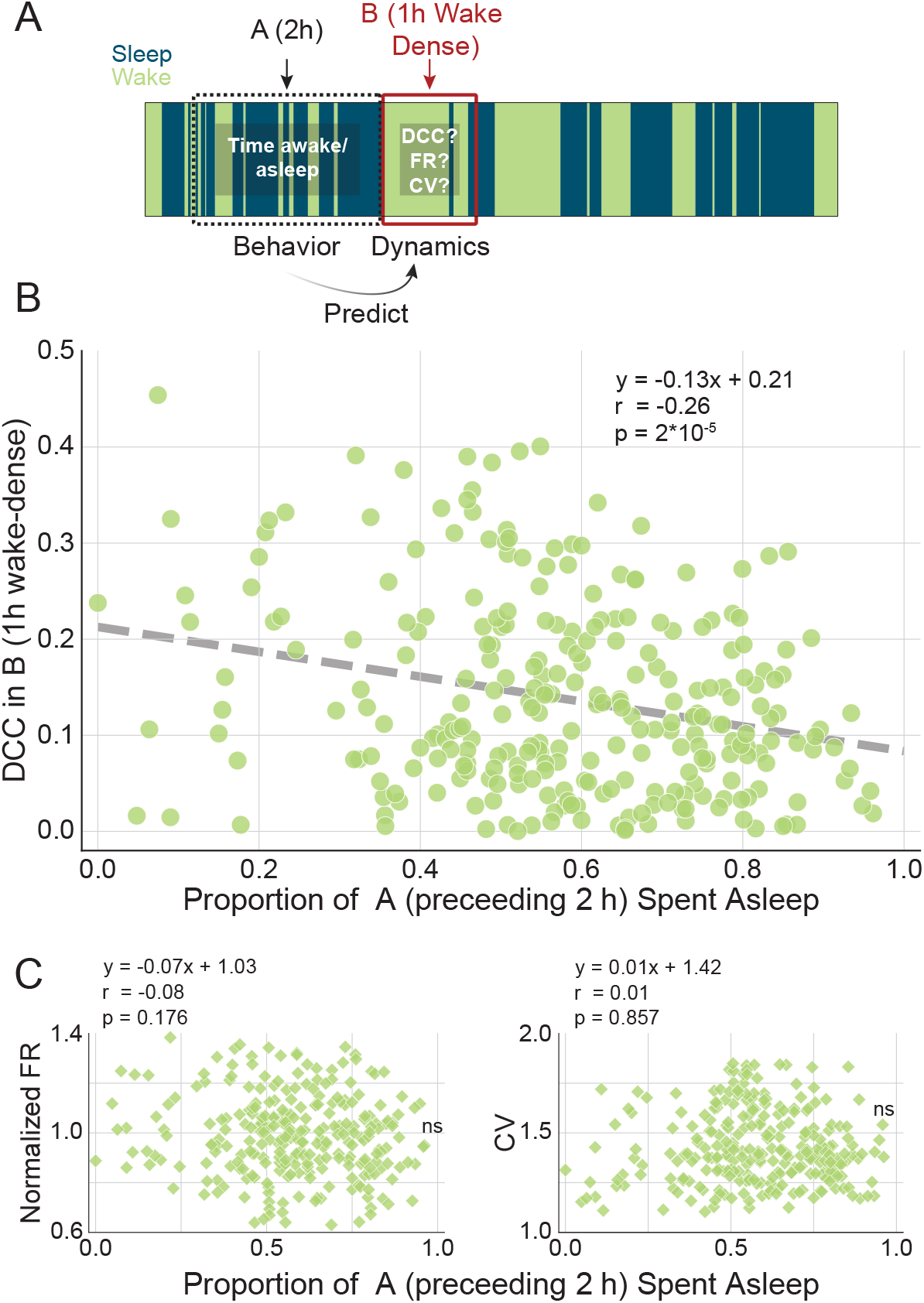
Prior behavior influences DCC within wake. (A) 1 h wake-dense epochs are identified (window B, red) in which features of neural activity are calculated. The amount of sleep in the preceding 2 h window (A, black) is calculated as the recent behavioral history. (B) Despite the ongoing effects of wake on DCC within each wake-dense block, there is a significant negative correlation in DCC during 1 h of wake and time spent asleep in the prior 2 h. (C) Analyzed in the same data, there is no relationship between sleep in the 2 h prior and normalized single unit firing rate (left) or coefficient of variation (right) in current wake-dense window.

## METHODS

All surgical techniques and experimental procedures were conducted in accordance with protocols approved by the Washington University in Saint Louis Institutional Animal Care and Use Committee (IACUC), following NIH guidelines for the care and use of research animals. Two female and six male Long-Evans rats (Charles River) were used in this study. Rats were 21 d postnatal (P21) at the time of surgery and P24 at the beginning of data collection.

### Animals and surgery

For electrode array implantation, rats were anesthetized with isoflurane (5% for induction, 2% for maintenance, mixed with air). Rats were head-fixed in a robotic stereotaxic instrument (Neurostar; Tubingen, Germany). The fur, skin, and periosteum covering the dorsal surface of the skull were removed, and the skull was then cleaned with hydrogen peroxide. Rats were administered slow-release Buprenorphine (0.1mg/kg) and Meloxicam (1 mg/kg) for pain relief. A craniotomy (1.8mm × 1.4mm) was drilled above the primary visual cortex (V1m) of the right hemisphere. To expose the surface of the brain, dura was resected by fine forceps or a 25G needle. Custom built microelectrode arrays were implanted into V1m at a rate of 5mm/minute using the stereotaxic robot and a custom vacuum holder. Arrays were bundles of 16 tetrodes (64 ch), composed of 12 μm ⌀ wire, cut to a length of 2-3 cm and bundled to create a flex cable leading back to a custom manufactured PCB (1.5 cm^2^ x 250 μm). Wires were soldered to the PCB (total weight of 300 μg). Implantation coordinates were 1.45/3.45/-1 (AP/ML/DV relative to Lambda and dura, in mm). The grounding wire was connected to a bone screw implanted in the skull over the left hemisphere. The implanted end of arrays was secured with dental cement, and, via the “flex cable”, headstage electronics (arrays and an adaptor; White Matter LLC, Seattle, WA) were assembled and housed in a custom 3D-printed enclosure. Total headstage weight including cement, bone screw, and enclosure was 2 g. Rats were administered Meloxicam (2 mg/kg) and Dexamethasone (1 mg/kg) at the end of surgery and over the next 2 d of recovery prior to connecting to the in vivo recording system. Animals were housed in an environmentally-enriched chamber with access to a litter mate through a perforated acrylic divider. Animals were kept on a 12:12 h light/dark cycle.

### In vivo recording and spike sorting

Freely behaving rats were attached to a custom-built tether with and in-line slip-ring (commutator). Neuronal signals were amplified, digitized, and sampled at 25,000 Hz using the eCube Server (White Matter LLC). Animal behavior was monitored with 15 or 30 fps video recording synchronized to the electrophysiological data with WatchTower (White Matter LLC). Recordings were conducted continuously for 10 – 14 d. Based on the recording stability and data quality across different animals, we used data of 4 - 10 d for future analysis. During recordings, in-person animal husbandry checks were conducted twice per day. In addition, live streams of data and video were monitored remotely using Open Ephys (open-ephys.org) and WatchTower (White Matter LLC).

Raw electrophysiological data were bandpass filtered between 500 and 7,500 Hz and thresholded for spike waveform extraction (mean ± 4 SD). Characteristics of isolated spikes were quantified for dimensionality reduction and then clustered by a modified version of SpikeInterface (Buccino et al., 2020) and MountainSort4 (Chung et al., 2017). Importantly, MountainSort4 clustering results were returned without curation, allowing for the examination of single-unit, multi-unit, and noise clusters. Single units were identified with an XGBoosted decision tree trained on tens of thousands of single units collected and manually scored in similar recordings. XGBoost results were evaluated for accuracy and curated by the experimenters, blind to the experimental specifics (i.e., spiking statistics, such as stability of waveform amplitude, cluster isolation, and refractory period contamination, were evaluated with no information about animal behavior or recording timeline). Data was clustered in blocks of 12 h. The mean number of single units extracted per block (12 h) was 35 +/- 7.9 (SEM).

### Sleep-wake scoring / polysomnography

Arousal states were first determined by a Random Forest (RF) model trained on thousands of hours of previously collected data. The RF model was initially trained on EEG, EMG, LFP, and the output of pose-estimation models run on acquired video (DeepLabCut; Mathis et al., 2018). Many features in this model are redundant and high-confidence state scores are readily acquired using broadband LFP and a sensitive measure of movement, similar to previous descriptions (Watson et al., 2016). Due to the ability of DeepLabCut to detect twitches and respiratory-related movements in addition to large locomotor movements, we combined this output with broadband LFP and the RF identified NREM, REM, active waking (locomotion), and quiet waking epochs with high confidence. This included brief intermediate states of semi-wake, i.e., microarousals, following REM epochs, as expected based on prior literature (dos Santos Lima et al., 2019; Miyawaki et al., 2017). Expert sleep-scorers manually examined all 240-336 h of data (alongside synchronized video) to confirm and/or correct RF output. All scoring (RF and manual curation) was conducted in 4s epochs using custom software (Python) (Figure S1A).

Briefly, low frequency LFP (0.1– 16 Hz) was extracted from five channels carrying single unit activity (an indication of high quality neurophysiological signal). The five channels’ LFP was then averaged. The corresponding spectrogram was generated for each hour of the recording. Highly sensitive pose-estimation was conducted on each frame of video recording using DeepLabCut (Mathis et al., 2018). LFP and movement data were synchronized and aligned. In V1 cortical recordings, NREM is characterized by high delta power (0.1-4 Hz), REM is characterized by low delta and high theta (6-8 Hz), as well as only cyclic respiratory movements. Quiet waking, in contrast, is characterized by low to intermediate delta power, intermediate theta power (lower than REM), and intermediate motor activity. Active waking could easily be identified along the single axis of motor output, as it is defined here by vigorous movement (e.g., grooming, eating) and/or locomotion.

### Neural avalanches analysis

Single unit spiking was discretized in time bins of 40ms. In other words, each neuron ‘s spike train was binarized into 0 (no activity) or 1 (has activity) for each bin. Activity across all recorded neurons (within an animal) was then integrated into a single vector (binarized activity was summed across neurons for each time bin). A network-activity threshold (either the 20th, 25th, 30th or 35th percentile) was determined on a per-animal basis for both sleep and wake. We used the 30th or 35th percentile as the threshold for avalanche extraction during waking, and the 20th or 25th percentile for sleep. Results were qualitatively robust across a range of thresholds, as previously described (Ma et al., 2019). However, we selected parameters that maximized the overall goodness of power law fitting within each state (for the entire recording). These parameters were selected prior to the calculation of the scaling relation (the basis of DCC) and then frozen for all subsequent analyses. For example, parameters for sleep state were calculated in the context of all sleep data in aggregate (i.e., prior to examining effects of sleep as a function of time, behavior, or environment). Once parameters were identified that returned the best power law fitting, they (the parameters) were frozen for subsequent analyses. In this way, we could ensure that we were robust to variance in the underlying power law exponents that might change in a state-dependent fashion. DCC is a measure of the scaling relation (Figure 1E, right) that can only be assessed if/when power law fitting is satisfied.

Individual avalanches contain successive time bins whose activity exceeds the threshold. Two descriptive measures of an avalanche include its size (S; the integrated network activity), and its duration (D; the elapsed time, or total number of bins). For systems operating in a near-critical regime, the probability (P) of observing avalanches of a given size or duration must follow a power law: P(S) ∼ S^α^ or P(D) ∼ D^β^. We fitted the avalanche size and duration distributions with truncated power laws using maximum likelihood estimation as described previously (Ma et al., 2019; Marshall et al., 2016). This generated two exponents (α and β). The goodness of the distribution fitting was evaluated via a power law test (Ma et al., 2019). Crucially, epochs of data that fail to pass power law tests are not eligible for subsequent analysis of a scaling relation. This occurs frequently in recordings during free behavior, and may be driven by noise or the mixture of multiple critical states whose individual power laws have quite different exponents. This phenomenon is called quasi-criticality, and has recently been described in computational models (Fosque et al., 2021).

In critical systems, the average size S⟩ of avalanches with a given duration must scale with the duration according to the equation: S⟩(T) ∼ T^η^. The empirically observed η was fitted by linear regression. If the network is operating at the critical regime, η is predicted by the exponent relationship equation η_predicted=(β-1)/(α-1). The absolute difference between the η_predicted and the fitted η is the “Deviation from Criticality Coefficient” (DCC), a quantitative measure of how close to criticality a system is operating (see Ma et al., 2019 for further details).

### Analysis of avalanches within behavioral states

Calculation of DCC requires significant numbers of avalanches to estimate underlying power laws. As a result, it is not feasible to extract DCC at the beginning of a short epoch and separately at the end of the same epoch. To understand whether DCC is progressively changed by time spent in sleep and wake, we divided recordings into 4 h windows. Each 4 h window contained multiple epochs of sleep and wake, although the total amount of sleep (or wake) varied dramatically (in some cases, as much as 80% of the 4 h was spent in one state or the other). We concatenated the waking spike times and extracted avalanches. The same was done for sleep. This allowed us to evaluate DCC within a each state as a function of the proportion of 4 h block spent in that state.

### Slow-wave Activity (SWA) quantification

SWA was quantified as described in Franken et al., 2001. Briefly, five channels (out of 64) with robust LFP activity were selected from each animal. For each of the five channels, broadband data (25 kHz) was downsampled to 200 Hz, and the five channels were averaged together to create a pseudo-EEG trace (EEG). The EEG signal was decomposed into different frequency bands using Fast Fourier Transform (SciPy; scipy.org), and the power spectral density was computed for the delta band (0.5 - 4 Hz) in 4 s windows. SWA was calculated per minute as the median delta power across 15 windows (i.e., 1 min of data). The median is robust to scattered noise. For prediction of future sleep/wake behavior, the mean SWA in 2 h windows was normalized across the whole recording in each animal, thus allowing pooling across animals (in other words, raw SWA values were not meaningful outside of the baseline context of each recording).

### Extended waking

We used gentle interventions to extend the length of every other naturally occurring waking epoch to at least 90 minutes. The protocol is similar to previously described methods (Hengen et al., 2016). Briefly, interventions included the introduction of novel objects, gentle stirring of bedding with a paintbrush, and tactile stimulation (brief tickling of whiskers or paws with a small paintbrush). Interventions were only deployed if and when animals showed signs of consolidated quiescent waking, as this is the entry point for NREM sleep. In many cases, after interventions ceased at 90 min, animals continued to engage in active exploration and play (Figure 4B). After each extended waking epoch, animals were allowed to sleep, wake, and sleep naturally before the next extended waking epoch. This paragram lasted for 24 hours and typically consisted of 7-10 extended waking cycles. During the entire 24 h, behavior states of rats were monitored via real-time LFP (Open Ephys) and video (WatchTower). Experimenters used night vision goggles (FVTGA, Baoding, China) for interventions during the dark cycle so as to avoid introducing a confound of externally driven visual input.

### Branching ratio

The branching ratio is a quantification of how activity grows or decays in a network. A branching ratio of less than 1 indicates that activity is progressively reduced over time, and a branching ratio of greater than 1 describes growth. Each of these regimes is unstable without further constraints. In critical systems, activity is self sustaining and neither explodes nor decays. This is achieved by a branching ratio of 1, which means that n spikes at time t will drive and average of n spike at time t+1. Previous theoretical work suggests that a branching ratio of slightly less than 1.0 (i.e., 0.97) is consistent with a critical system in the context of external input (Ma et al., 2019). To quantify the branching ratio in recorded spiking, we employed an open toolbox that is particularly powerful due to its robustness against subsampling (Spitzner et al., 2021); only a small subset of the neurons in the entire visual cortex are recorded here.

Details about the toolbox can be found in Spitzner et al., 2021. Briefly, the toolbox uses a Multistep Regression Estimator to calculate the branching ratio in three principal steps. First, spike times of neural ensembles are binarized into 4ms time bins, and network activity is calculated as the sum of all binarized activity within the time bin. Second, correlation coefficients are calculated between network activity at time bins t and t+k (linear regression), with different time lags k from 1 to kmax. These correlation coefficients of different time lags typically feature an exponentially decaying autocorrelation function. Third, the curve of correlation coefficients is fit with one of a variety of fitting functions embedded in the toolbox. In this study, we found that the intermittent off states that characterize NREM sleep are best navigated by the “complex” fitting function, while the more continuous spiking that characterizes waking is captured by the “exponential_offset” function. These two options yielded the best goodness of fit across all animals. We used a kmax of 500 in all instances.

### Analysis of wake-dense and sleep-dense blocks

To analyze wake-dense or sleep-dense blocks for each animal, a 1-hour window was slid over the data and each non-overlapping block with at least 66% time spent awake or asleep was identified (using the find_peaks function in the Scipy package). DCC was calculated in each of these windows, and then correlated with the percentage of sleep in the 2 h immediately preceding the start time of the sleep- or wake-dense window.

### Machine learning models for sleep prediction

We employed logistic regression models and XGBoosted decision trees models to predict sleep- and wake-dense future 1 h windows based on either single features (logistic regression) or combinations of features (XGBoost) measured in the preceding 2 h windows. 80% of the dataset was used for training models and 20% of the dataset was withheld for testing (prediction accuracy is based on the test set). Data from all 8 rats were included in each training and testing set. Parameters of the XGBoost model were determined by hyperparameter tuning to optimize the prediction accuracy and avoid overfitting of the training dataset. Because the splits selected for train and test could, purely due to chance, lead to artificially high or low accuracy, each XGBoost model was run 200 times and each logistic regression model was run 50 times (XGBoost is more sensitive to data splitting as it contains more features and is nonlinear). In each iteration, a new randomly train/test split was chosen. The accuracy of the model is defined as the average prediction accuracy across all iterations. To measure feature impotence, weights of individual features were extracted from and averaged across the 200 XGboost models trained on all features (behavioral history, normalized firing rate, CV of ISI, DCC, and ZT).

### Probe Localization

After recordings were completed, rats were euthanized and perfused with 4% formaldehyde (PFA). Brains were extracted and fixed for 24 h at 4°C in PFA. Brains were then transferred to 30% sucrose solution and stored at 4°C for 48 h. Brains were sectioned at 60 μm on a cryostat. Sections were rinsed in PBS prior to being mounting on charged slides (SuperFrost Plus, Fisher) and stained with cresyl violet using a standard protocol. Tetrode tracks were identified under a microscope and confirmed in serial sections.

### Statistics

For visualization, data are reported as mean ± SEM for 8 animals, unless otherwise noted. For statistical comparisons, a linear mixed effect (LME) model is used with a significance level of p<0.05 unless otherwise noted (Python Statsmodels Module). Where noted, a T-test (for normally distributed data) or Mann-Whitney test (for non-normally distributed data) are employed. One-way single-factor ANOVA followed by a post hoc Tukey test is used for comparing multiple groups. The Kolmogorov-Smirnov test is used to compare cumulative distributions.

